# Obstacle avoidance in aerial pursuit

**DOI:** 10.1101/2023.01.23.525170

**Authors:** Caroline H. Brighton, James A. Kempton, Lydia A. France, Marco KleinHeerenbrink, Sofia Miñano, Graham K. Taylor

## Abstract

Collision avoidance [1–4] and target pursuit [5–8] are challenging flight behaviors for any animal or autonomous vehicle, but their interaction is even more so [9–11]. For predators adapted to hunting in clutter, the demands of these two tasks may conflict, requiring effective reconciliation to avoid a hazardous collision or loss of target. Technical approaches to obstacle avoidance rely mainly on path-planning algorithms [12], but these are unlikely to be effective during closed-loop pursuit of a maneuvering target, so collision avoidance must instead be implemented reactively during prey pursuit. For example, the pursuit-avoidance behavior of predatory flies has been successfully modelled by combining feedback on target motion with feedback on obstacle looming [13]. It is unclear, however, whether this mechanism will generalize to complex environments with many looming obstacles, and it remains unknown how aerial predators reconcile the conflict between obstacle avoidance and prey pursuit in clutter. Here we use high-speed motion capture data to show how Harris’ hawks *Parabuteo unicinctus* avoid collisions by making open-loop steering corrections during closed-loop pursuit. We find that hawks combine continuous feedback on target motion with a discrete feedforward steering correction aimed at clearing an upcoming obstacle as closely as possible at maximum span. By biasing the hawk’s flight direction, this guidance law provides an effective means of prioritizing obstacle avoidance whilst remaining locked-on to the target. We anticipate that a similar mechanism may be used in terrestrial and aquatic pursuit. The same biased guidance law could be used for obstacle avoidance in drones designed to intercept other drones in clutter, or in drones using closed-loop guidance to navigate between fixed waypoints in urban environments.

---

Pursuing prey through clutter is a complex and risky activity requiring the integration of distinct guidance subsystems for obstacle avoidance and target pursuit. Harris’ hawks offer an excellent model system for studying this problem, because their pursuit behavior has been well characterized in the open [11], but their hunting strategy involves making short flights after terrestrial prey in habitat clutter [14]. Previous work has found that their unobstructed pursuit trajectories are well modelled by assuming that turning is commanded at an angular rate:

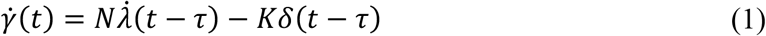

where *N, K* and *τ* are fitted constants, where 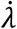 is the angular rate of the line-of-sight from the pursuer to the target, where *δ* is the signed deviation angle between the pursuer’s flight direction and its line-of-sight to the target, and where *t* is time [11]. Here, we use a high-speed motion capture system to reconstruct the flight trajectories of *N* = 4 hawks chasing a lure towed along an unpredictable path about a series of pulleys in a large hall with or without obstacles (Video 1). We then simulate these data computationally using several alternative models of the guidance dynamics, which we use for hypothesis testing.

## Experimental design

We used two rows of hanging ropes as obstacles: the first forming a dense clump that the bird had to fly around, and the second simulating a row of trees that the bird had to fly between (Fig. 1A,B). The full dataset contains four subsets: (i) a set of n=128 obstacle-free training flights collected over 8 days; (ii) a set of n=16 obstacle familiarization flights collected the next day; then (iii) a set of n=106 obstacle-free test flights; and (iv) a set of n=154 obstacle test flights; where (iii) and (iv) were collected over 15 days on which the presence or absence of obstacles was randomized (see Methods). The n=106 obstacle-free test flights are reported and included in an analysis of unobstructed pursuit elsewhere [15], but the flights with obstacles are described here for the first time.

**Figure 1.**
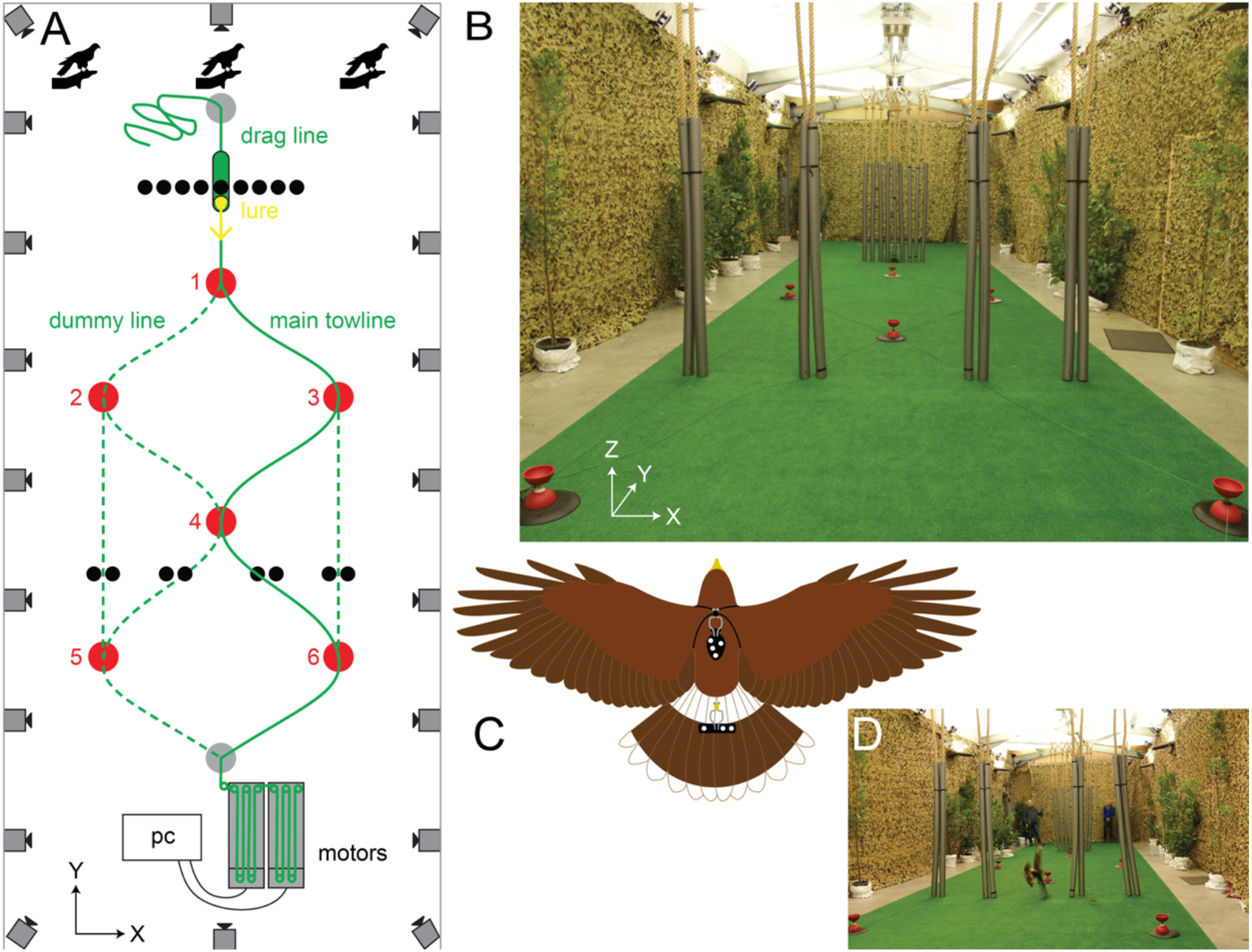
Experimental setup. **(A)** Overhead schematic of flight hall. Each of the N=4 Harris’ hawks flew from one of three alternative starting positions (bird icons), chasing a food lure (yellow arrow) pulled by a pair of linear motors on a towline (solid green line) with a trailing drag line that ran around 3 or 4 out of 6 pulleys (red dots). Dummy towlines (dashed green lines) were laid around the remaining pulleys, to prevent the bird from anticipating which of the 6 alternative paths the lure would follow. The hawk and lure were tracked by 20 motion capture cameras positioned around the room (camera icons). Ropes (black dots) were hung as obstacles in the configuration shown for the test flights with obstacles. **(B)** View of experimental set-up looking from the linear motors back towards the starting positions of the bird and lure; note the diffuse overhead lighting provided by bouncing light from the 8 LED up-lights positioned around the walls. Shrubs and trees were placed down the sides of the room to provide visual contrast and discourage flight outside of the central test area. **(C)** Schematic showing the marker templates (black patches) worn on the back and tail of the hawk, together with their attached retroreflective markers (white dots). **(D)** Cropped frame from Video 1, showing example flight.

## Model validation

We begin by using our new sample of n=128 obstacle-free training flights to validate the mixed guidance law (Eq. 1). We match the hawk’s simulated speed to its measured speed and use Eq. 1 to model its horizontal turning behavior, taking the measured trajectory of the lure as a given, and matching the initial conditions of each simulation to the measured data. We define the prediction error of the simulation, *ε*(*t*), as the distance between the measured and simulated trajectories, which we summarize by reporting the mean prediction error 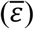 for each flight, and its median 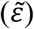 over all the flights within a subset. Simulating the obstacle-free training flights at the published [11] parameter settings of *N* = 0.7, *K* = 1.2 *s*^-1^ and *τ* = 0.09 *s* typically resulted in a low mean prediction error, with a median value of 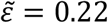 m over the n=128 flights (95% CI: 0.20, 0.28 m). By comparison, the median over the independent data set of n=50 obstacle-free flights to which Eq. 1 was originally fitted was 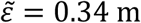 (95% CI: 0.24, 0.53 m). Eq. 1 therefore models our sample of n=128 obstacle-free training flights at least as well as the sample of n=50 outdoor flights to which it was fitted, validating its suitability as a model of unobstructed pursuit behavior in Harris’ hawks. Because Eq. 1 feeds back the deviation angle *δ*, it produces a characteristic tail-chasing behavior that is hypothesised to promote implicit collision avoidance when chasing a target that is itself weaving between obstacles [11]. The lure travelled through the gaps between obstacles on the n=16 obstacle familiarization flights, so we tested this hypothesis by using Eq. 1 to simulate these flights at the parameter settings above. Although the model does not always predict the hawk’s turning behavior closely at the point of capture, it predicts the earlier sections of each flight well, following the lure through the gaps between obstacles (Fig. S1A). The target pursuit subsystem that Eq. 1 describes is therefore capable of producing a safe path through clutter when chasing a target that itself passes safely between obstacles.

## Model refinement

We next refined the parameters of the mixed guidance law (Eq. 1) by re-fitting these to the n=260 test flights that we recorded. For direct comparability with the results of our modelling using the original mixed guidance law [11], all of our simulations begin from 0.09 s after the start of each recording, allowing the fitting of a sensorimotor delay of *τ* ≤ 0.09 s. We began by fitting separate models to the test flights with and without obstacles, finding the guidance parameter settings that minimized the median of the mean prediction error, 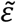, over each subset of flights (see Materials and Methods). However, as the optimized parameters were similar for each subset (*N* = 0.75, *K* = 1.15 s^-1^ and *τ* = 0.005 s for the n=106 obstacle-free test flights; *N* = 0.75, *K* = 1.15 s^-1^ and *τ* = 0.015 s for the n=154 obstacle test flights), and were close to those fitted in previous work [11], we re-fitted the model to the union of the test flights with and without obstacles. Because flights with obstacles are overrepresented in this sample relative to flights without obstacles, we used a subsampling procedure in which we randomly subsampled 80 flights without replacement from each subset and identified the parameter settings that minimized 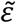 over that subsample (see Materials and Methods). We repeated this sampling experiment 100,000 times and took the median of the best-fitting parameter settings as our refined model. The goodness of fit of this model with refined parameter settings of *N* = 0.75, *K* = 1.25 s^-1^ and τ = 0.010 s was similar for the n=106 obstacle-free test flights (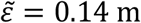; 95% CI: 0.12, 0.19 m; Fig. S1B) and the n=154 obstacle test flights (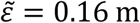; 95% CI: 0.14, 0.21 m; Fig. 2, S1C). Moreover, it performed marginally better on the validation data from the n=128 obstacle-free training flights (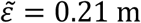; 95% CI: 0.17, 0.26 m) than the original version of the mixed guidance law [11]. We therefore take this refined mixed guidance law as our best-supported model of the target pursuit subsystem of Harris’ hawks.

**Figure 2.**
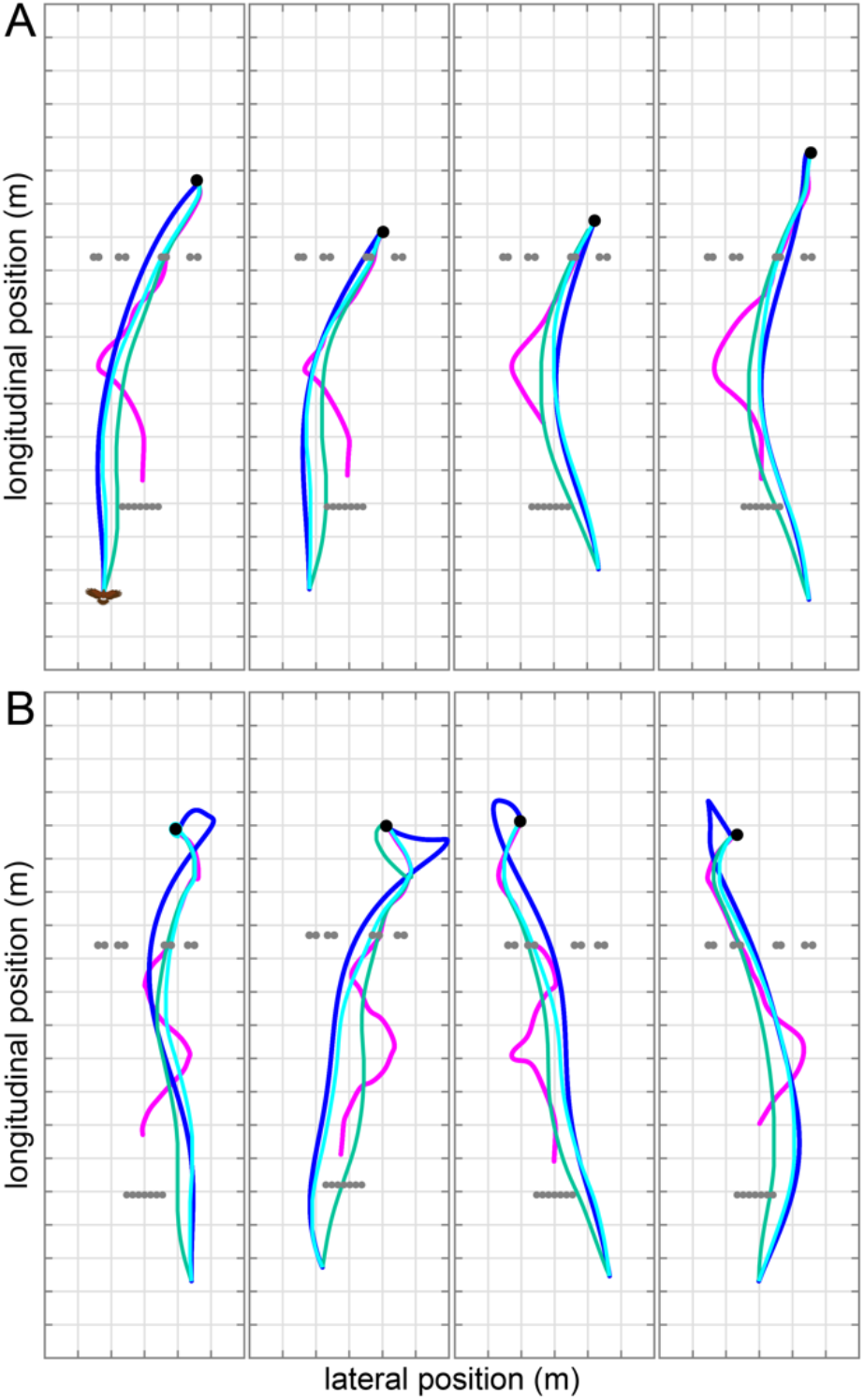
Effect of measured take-off bias on pursuit simulations. Each panel represents a single obstacle test flight and plots the hawk’s measured flight trajectory (blue) in pursuit of the lure (magenta) up to the point of capture (black dot). Hanging rope obstacles are plotted as grey dots. The measured data are compared to simulations of the hawk’s trajectory (cyan) under the refined mixed guidance law (Eq. 1), with best-fitting parameters *N* = 0.75, *K* = 1.25 s^-1^ and *τ* = 0.01 s, with the initial deviation angle *δ*_0_ matched to the value measured. These are compared to simulations with the initial deviation angle set to *δ*_0_ = 0, simulating take-off directly towards the lure (green). **(A)** The four flights > 9 m in length having the lowest mean prediction error relative to distance flown: whilst there are shorter flights with lower absolute mean prediction error 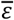, these are among the most closely fitted flights. **(B)** The four longest flights recorded: these are not amongst the most closely fitted flights, with all four having a mean absolute prediction error 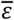 significantly higher than the median 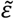. Grid spacing: 1 m.

## Take-off direction is biased to avoid obstacles

The refined mixed guidance law usually predicted a collision-free path around the first row of obstacles (Fig. S1C), which reflects the fact that our simulations are initialized using the bird’s measured take-off velocity. Hence, if a hawk sets its take-off direction to avoid the first set of obstacles, then the resulting bias in the initial value of its deviation angle *δ* will be embedded in its simulated pursuit behavior. We tested this prediction by comparing the distribution of the initial deviation angle, *δ*_0_, measured between the hawk’s flight velocity and its line-of-sight to the lure at the start of the simulation, for the different test flight subsets (Fig. 3). Whereas the distribution of *δ*_0_ was unimodal with a mode at *δ*_0_ ≈ 0° for the test flights without obstacles, it was bimodal with modes at *δ*_0_ ≈ ±20° for the test flights with obstacles (Fig. 3A). Accordingly, the median absolute initial deviation angle (Fig. 3B) was larger for the test flights with obstacles (21.2°; 95% CI: 19.8°, 23.8°; n=154 flights) than for those without (11.7°; 95% CI: 9.1°, 13.9°; n=103 flights; see Fig. 3 legend for details of 3 exclusions). Hence, whereas the hawks took off towards the lure when there were no obstacles present, they biased their take-off away from any obstacle that was blocking their path to the lure. We next tested whether this observed bias in take-off direction was necessary and sufficient to ensure that the hawk’s target pursuit subsystem would produce a safe path around the first obstacle. We checked this by re-running the simulations for the test flights with obstacles under the refined mixed guidance law, having set the initial deviation angle as *δ*_0_ = 0 (i.e., having set the simulation to take off directly towards the lure, despite the presence of an obstacle blocking the way). These simulations often produced a collision with the first obstacle, even when no collision had been predicted with *δ*_0_ set to the value that we observed (Fig. 2). It follows that the hawks’ observed bias in take-off direction was both necessary and sufficient to cause their target pursuit subsystem (Eq. 1) to produce a safe path around the first obstacle.

**Figure 3.**
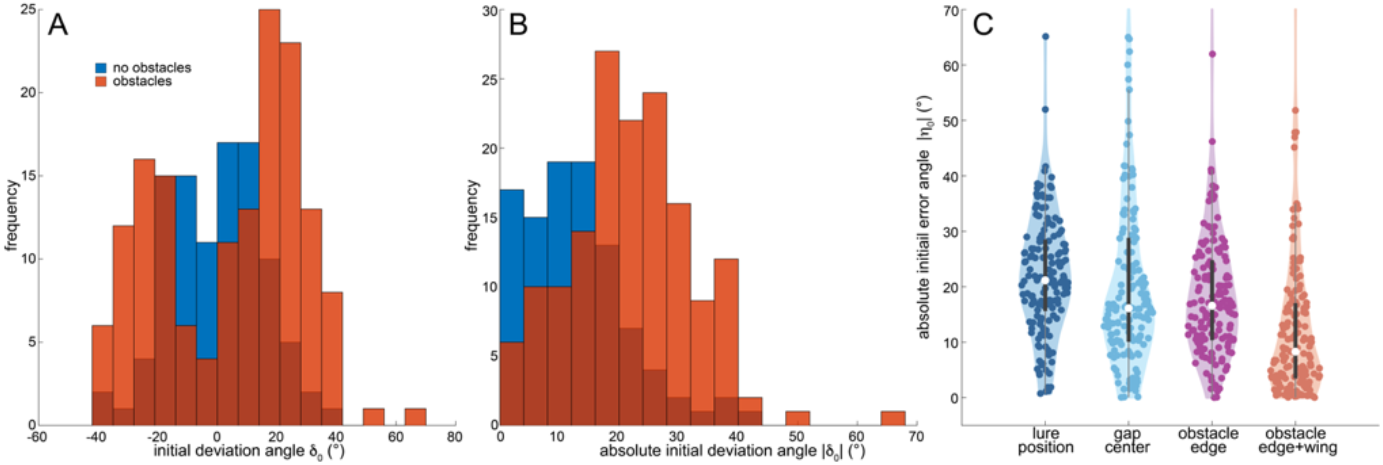
Distribution of take-off bias. **(A)** Histogram of the initial deviation angle, *δ*_0_, defined as the angle between the hawk’s flight direction and its line-of-sight to the lure, sampled at the time *t* = 0 from which the guidance simulations began. **(B)** Histogram of the absolute initial deviation angle, |*δ*_0_|. **(C)** Violin plots of the absolute initial error angle, | *η*_0_|, defined as the angle between the hawk’s initial flight direction and its initial line-of-sight to the target defined on the *x-* axis. In the special case that the target is the lure, |*η*_0_| ≡ |*δ*_0_|. The three alternative target definitions include: (i) the nearest edge of the first obstacle; (ii) the center of the gap between this and the wall; (iii) an intermediate position approximately one wing length (0.5 m) into the gap from the edge of the obstacle. Data are shown for all n=154 obstacle test flights, and for n=103 obstacle-free test flights, having dropped all 3 flights on which the hawk had already travelled beyond the location of the first obstacle by the point at which the guidance simulations began.

## Take-off bias minimizes obstacle clearance at maximum span

How did the hawks select an appropriate take-off bias? Previous research on obstacle avoidance has found that domestic pigeons *Columba livia domestica* target the centers of gaps between obstacles [2, 16], and that Harris’ hawks look at the nearest edge of obstacles they are avoiding [17]. We therefore hypothesized that the hawks took off by aiming at either the nearest edge of the obstacle or the midpoint of the gap between the obstacle and the wall. We tested this by calculating the initial error angle, *η*_0_, between the hypothesized take-off aim and the direction of the hawk’s flight and compared this to the equivalent error angle for the lure (i.e., the initial deviation angle *δ*_0_). The median absolute initial error angle was smaller (Fig. 3C) when the hawk was assumed to have aimed its take-off at either the obstacle edge (median |*η*_0_|: 16.6°; 95% CI: 15.0, 18.6) or the gap center (median |*η*_0_|: 16.1°; 95% CI: 14.4, 17.6) rather than the lure (median |*δ*_0_|: 21.2°; 95% CI: 19.8°, 23.8°). However, the initial error angle was smaller again if the hawk was assumed to have aimed for a clearance of approximately one wing length (0.5 m) from the obstacle edge (median |*η*_0_|: 8.3°; 95% CI: 6.2°, 10.7°), with the median absolute error angle, 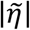, reaching a global minimum of 5° assuming a targeted clearance of 0.6 m on approach to the first obstacle (Fig. 4A,C). This strategy makes sense, because aiming at the edge of an obstacle leaves no clearance and aiming at the center of a gap leaves more clearance than is necessary for a gap larger than the wings’ span. We conclude that the hawks biased their take-off direction to turn tightly around the obstacle without having to close their wings, thereby reconciling any initial conflict between obstacle avoidance and target pursuit without limiting their control authority.

**Figure 4.**
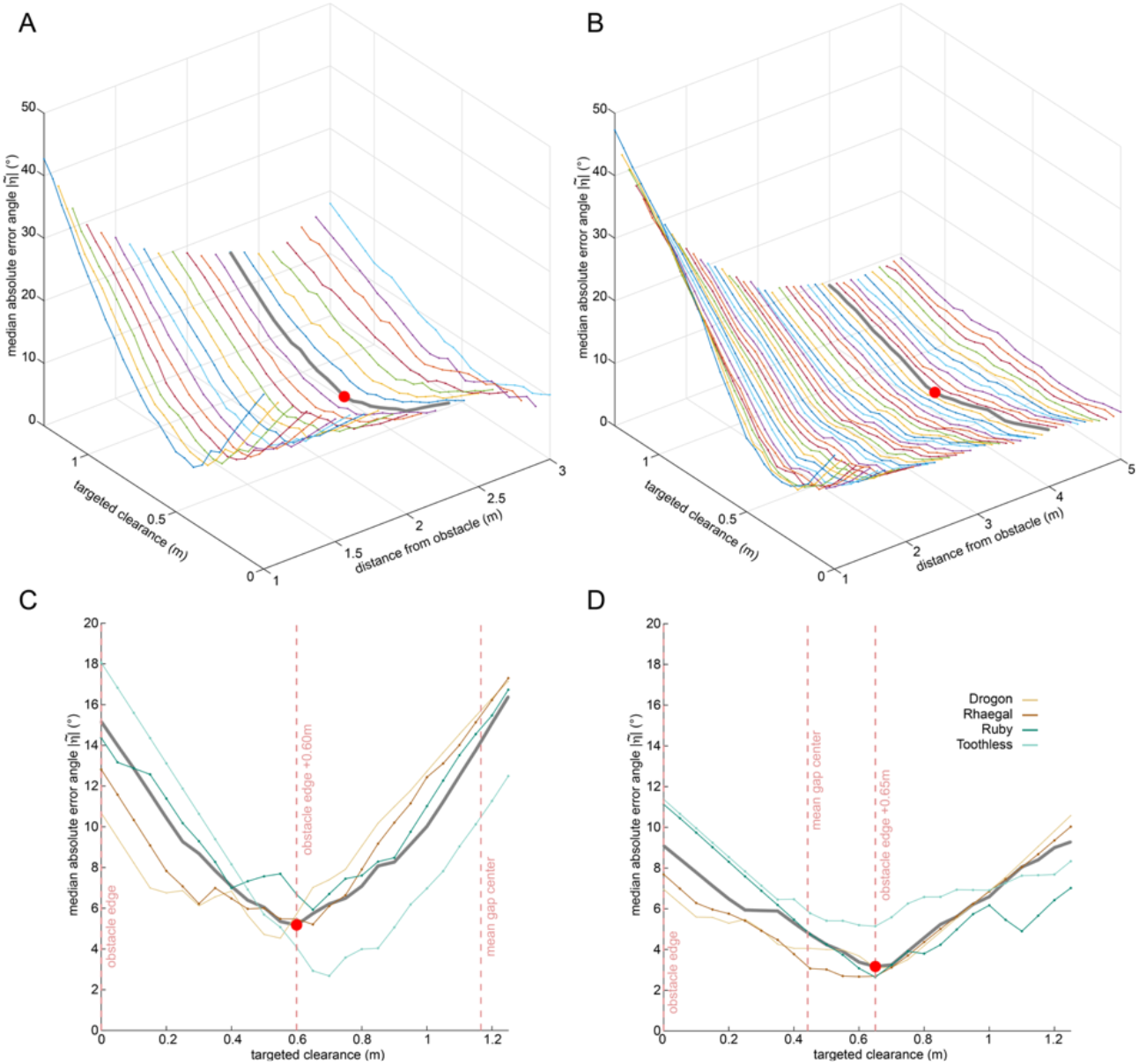
Error angle as a function of targeted clearance. Plots of the median absolute error angle, 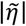, where the error angle *η* is defined as the angle between the hawk’s flight direction and its line-of-sight to the clearance, conditional upon the clearance being targeted. Data are shown for the n=111 obstacle test flights on which the hawk intercepted the target after passing the second obstacle. **(A,B)** Median absolute error angle 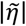 as a function of targeted clearance from: (A) the first obstacle; and (B) the second obstacle, plotted at a range of different distances from the obstacle. The global minimum (red dot) is reached at 2.2 m from the first obstacle (grey line), shortly after take-off, and at 4.0 m from the second obstacle (thick grey line). **(C,D)** Median absolute error angle 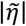 as a function of targeted clearance from: (C) the first obstacle; and (D) the second obstacle, plotted for the specific distance at which the global minimum is reached (thick grey line). The colored lines plot the same quantities for the subset of flights from each individual bird. Red dashed lines denote the locations of the targeted clearances referred to in the main text; note that the exact position of the gap center varies between trials owing to variation in the placement of the obstacles and is therefore summarized by its mean position across trials.

## Mid-course steering bias minimizes obstacle clearance at maximum span

The hawks’ initial bias in take-off direction explains how they avoided colliding with the first obstacle whilst chasing the target, but not how they avoided colliding with the second (see Fig. 2). We therefore looked for evidence of mid-course steering correction by comparing the time history of the median prediction error 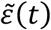 under the refined mixed guidance law for the n=154 test flights with obstacles and the n=106 test flights without (Fig. 5). Because the initial conditions of each simulation were matched to those we had measured, 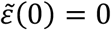 at the start of each flight. Thereafter, the simulations deviate from the measured trajectories, but do so to a greater extent when obstacles are present (Fig. S1B,C). This difference is consistent with the hypothesis that the hawks made mid-course steering corrections for obstacle avoidance that the simulations under Eq. 1 alone do not capture. Moreover, the median prediction error 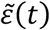 peaks at the times the hawks passed the first and second obstacles but does not peak at those times for the test flights without obstacles (Fig. 5). The hawks therefore deviated most strongly from the trajectory commanded by their target pursuit subsystem as they negotiated obstacles, providing clear evidence of mid-course steering correction to avoid these. Given the biased take-off mechanism that we have already identified, we hypothesize that mid-course steering correction will likewise involve aiming for a clearance of approximately one wing length from any obstacle blocking the path to the target. To test this hypothesis, we repeated the error angle analysis that we had undertaken for the first obstacle (Fig. 4A,C), computing how the error angle, *η*, varied on approach to the second obstacle in relation to the bird’s assumed steering aim (Fig. 4B,D). Consistent with the results for the first obstacle (Fig. 4A,C), we found that the median absolute error angle, 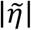, reached a global minimum of 3° when the hawks were assumed to aim for a clearance of 0.65 m from the obstacle (Fig. 4B,D). This minimum was reached 4 m from the second row of obstacles (Fig. 4B), so the hawks appear to have made a mid-course steering correction by the time they were within 4 m of the second obstacle.

**Figure 5.**
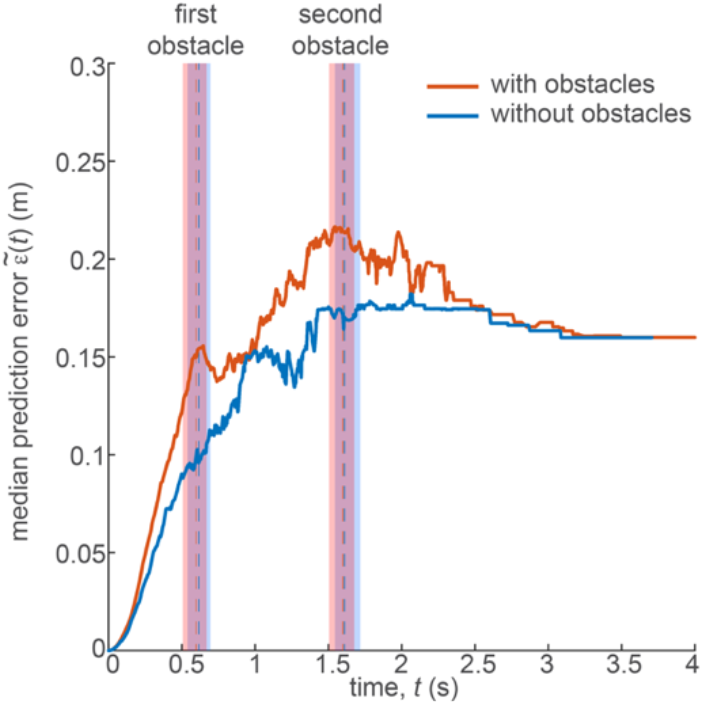
Median prediction error against time. Median prediction error 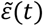 between the measured flight trajectories and those simulated under the refined mixed guidance law (Eq. 1), with best-fitting parameters *N* = 0.75, *K* = 1.25 s^-1^ and *τ* = 0.01 s. Because the initial conditions of each simulation were matched to those we had measured, 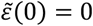 by definition. The simulations deviate from the measured trajectories over time but do so to a greater extent on the n=154 obstacle test flights (orange) than on the n=106 obstacle-free test flights (blue). The dashed lines and vertical bars denote the median and interquartile range, respectively, of the times at which the hawks passed the locations of the first and second obstacles. Note that the median prediction error peaks at these times for the test flights with obstacles (orange) but not for the test flights without obstacles (blue), providing evidence of mid-course steering correction to avoid them.

## Mid-course steering bias is applied in feedforward fashion

Avoiding obstacles by aiming at a clearance lends itself well to open-loop steering correction, which is the simplest way in which the intermittent demands of obstacle avoidance may be combined with the continuous demands of target pursuit. Under this hypothesis, a feedforward bias command would be issued at some threshold distance (or time to collision) from an upcoming obstacle, aimed at perturbing the pursuer’s deviation angle *δ* such that the continuation of its pursuit begins with the pursuer heading for a clearance of approximately one wing length from the near edge of the obstacle. In contrast, previous studies of obstacle avoidance in pigeons [2, 16] have modelled obstacle-avoidance as a closed-loop steering behavior, treating the gap between obstacles as a goal that the bird uses feedback to steer toward. Superposing the resulting steering command with that of the target pursuit subsystem would result in a composite steering command representing a continuous compromise between target pursuit and obstacle avoidance. Provided the tuning of the guidance parameters is similar for both subsystems, their composite steering command is economically modelled by redefining the target of Eq. 1 as the point midway between the lure and the gap. This simple approach ensures that we continue to fit only three guidance parameters in the analysis of closed loop steering that follows.

To test whether there was evidence for closed-loop steering to avoid obstacles, we re-fitted the parameters of the mixed guidance law to the n=111 obstacle test flights on which the hawk intercepted the target after passing the second obstacle, redefining the target of Eq. 1 as the point midway between the lure and a clearance of 0.6 m from the near-edge of the second obstacle. As before, we matched the initial conditions of the simulations to those we had measured. For comparison, we also fitted the simulations treating either the lure or the assumed clearance from the obstacle as the target of Eq. 1. In each case, we only fitted the simulations as far as the second row of obstacles, to avoid the need to redefine the target at this point. The prediction error was smallest for the simulations treating the lure as the target (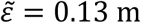; 95% CI: 0.12, 0.16 m), largest for the simulations treating the assumed 0.6 m clearance as the target (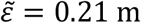; 95% CI: 0.20, 0.26 m), and intermediate for the model targeting the point midway between them (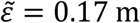; 95% CI: 0.16, 0.19 m). This analysis therefore provides no evidence of closed-loop steering towards the gap between the obstacles, although it does not exclude the possibility that some other mechanism of closed-loop obstacle avoidance was in operation.

We are left with the hypothesis that Harris’ hawks pursue targets through clutter under the mixed guidance law identified above, but that they avoid upcoming obstacles using feedforward steering commands. To model this behavior we: (i) inherited the parameters of the refined mixed guidance law that we had fitted already (i.e *N* = 0.75, *K* = 1.25 s^-1^ and *τ* = 0.01 s); (ii) prescribed the initial conditions by aiming take-off for a clearance of 0.6 m from the near-edge of the first obstacle; and (iii) added a discrete change in flight direction 4 m ahead of the second obstacle, aiming this for a clearance of 0.6 m from the near-edge of the obstacle closest to the hawk’s flight direction. In cases where the obstacles were spaced less than 1.2 m apart, such that aiming for a clearance of 0.6 m from one would have brought the bird closer than 0.6 m to the other, we aimed this change in flight direction at the center of the gap between them. We used this model to simulate the n=111 obstacle test flights on which the hawk intercepted the target after passing the second obstacle (Fig. 6), and found that it fitted these data marginally better (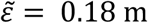; CI: 0.14, 0.22 m) than the refined mixed guidance law with initial conditions matched to those we had measured (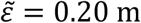; CI: 0.15, 0.25 m). Combining a feedforward command in response to upcoming obstacles with a feedback command in response to target motion therefore provides an effective means of prioritizing obstacle avoidance whilst remaining locked-on to the target, and closely explains our data.

**Figure 6.**
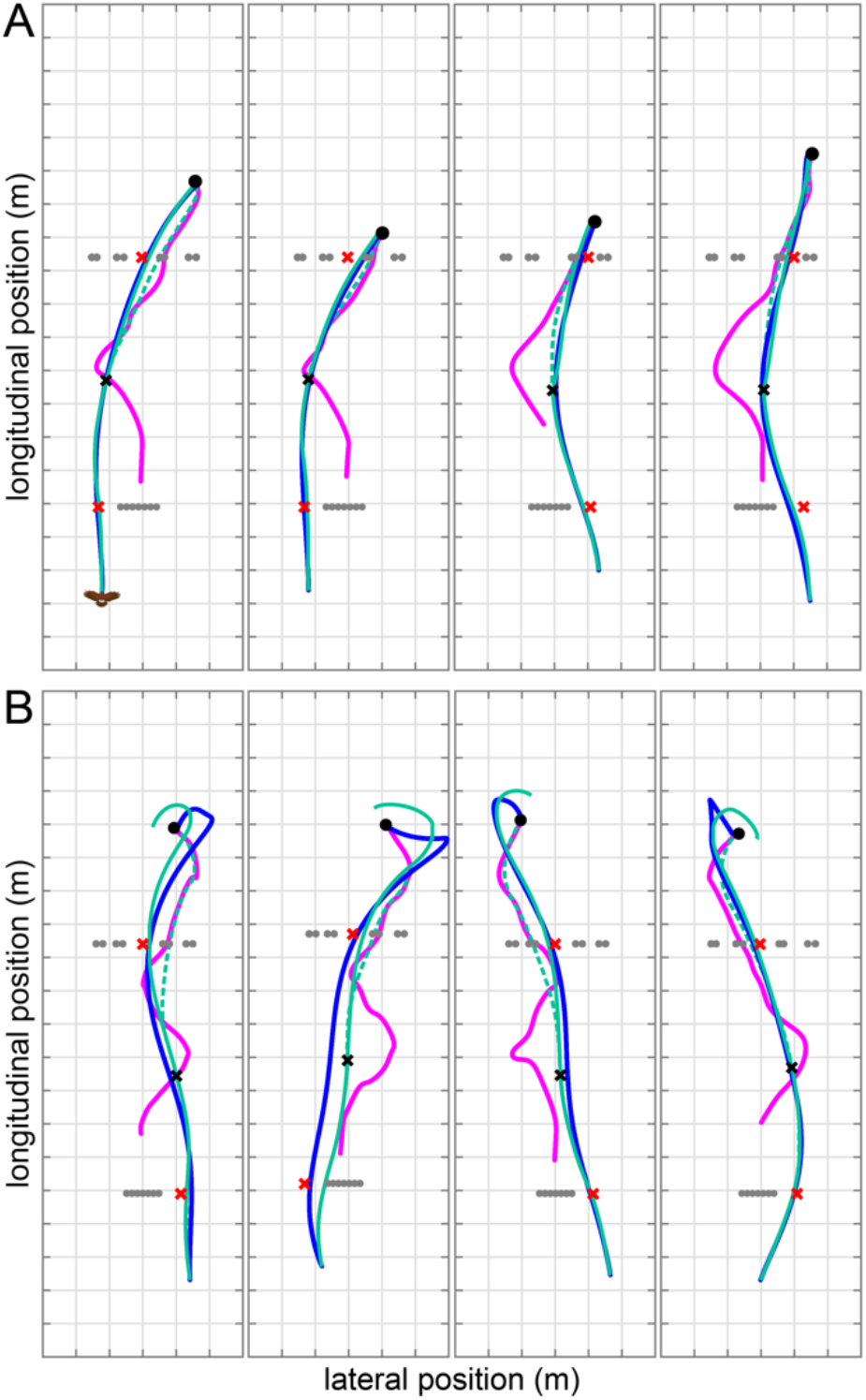
Simulations of closed-loop pursuit with feedforward obstacle avoidance. Each panel represents a single obstacle test flight and plots the hawk’s measured flight trajectory (blue) in pursuit of the lure (magenta) up to the point of capture (black dot). Hanging rope obstacles are plotted as grey dots. The measured data are compared to simulations of the hawk’s trajectory (green) under the refined mixed guidance law (Eq. 1), with best-fitting parameters *N* = 0.75, *K* = 1.25 s^-1^ and *τ* = 0.010 s, and with a discrete bias command applied once upon take-off and once when the hawk reaches 4.0 m from the second obstacle (black cross). Each bias command is implemented as an instantaneous steering correction targeting a clearance of 0.6 m from the nearest edge of an upcoming obstacle (red crosses). In cases where the gap between obstacles was <1.2 m, the steering correction was assumed to target the center of the gap. The dashed green line plots the continuation of the simulation without the second steering correction applied, showing the effect of its application on obstacle avoidance. **(A)** The four flights > 9 m in length having the lowest mean prediction error relative to distance flown: whilst there are shorter flights with lower absolute mean prediction error 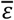, these are among the most closely fitted flights. **(B)** The four longest flights recorded: these are not necessarily amongst the most closely fitted flights, with three having a mean absolute prediction error 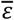 significantly higher than the median 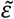. Grid spacing: 1 m.

## Target overshoot and residual collision risk

Unsurprisingly, our simulations do not capture every detail of the hawks’ turning behavior. In particular, the longest test-flight trajectories that we recorded ended with the hawk overshooting the lure and making a hairpin turn to catch it. This behavior was not captured by the refined mixed guidance law alone (Fig. S1B,C), yet perturbing the trajectory commanded by the target pursuit subsystem by adding a feedforward steering correction to avoid the second obstacle often caused the simulations to overshoot the lure in a more lifelike manner (Fig. 6A). The fact that a similar overshoot was also observed on the test flights without obstacles may suggest that the real birds were unable to generate an accurate steering command (e.g. because of sensor error), or were unable to meet this steering demand (e.g. because of physical constraint). It is also possible that this overshoot was adaptive, reflecting an aspect of the control of the final strike maneuver that our guidance simulations do not capture. Finally, although the hawks steered to avoid the obstacles we presented, the compliant nature of their wings and the ropes used as obstacles meant they could tolerate occasional collisions, like those they would experience when brushing past vegetation in their natural environment. Our feedforward model of obstacle avoidance led to a residual collision risk of 7% across the first and second obstacles, which closely matches the observed collision rate of 6%.

## Biased guidance as a biological strategy for pursuit-avoidance

Formally, we have evidence for biased guidance of obstructed pursuit in Harris’ hawks, with turning commanded at an angular rate:

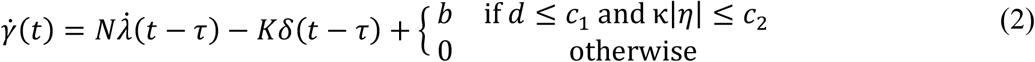

Here, *b* is a bias command, *d* is the distance to an upcoming obstacle, and *η* is the signed error angle between the pursuer’s flight direction and its line-of-sight to the near edge of the obstacle. In the case of our Harris’ hawks, we have the fitted guidance parameters *N* = 0.75, *K* = 1.25 s^-1^, and τ = 0.01 s, with *c*_1_ = 4 m and *c*_2_ = sin^-1^ (0.6/d) as constants defining the threshold distance and error angle at which obstacle avoidance is triggered. The variable κ takes the value κ = —1 if the pursuer is on a direct collision course with the obstacle, with κ = 1 otherwise, such that *c*_2_ defines the error angle tolerance with which obstacles are avoided. Our implementation of Eq. 2 in Fig. 6A assumes that the bias command is applied in open-loop, over a short time step of duration Δ*t*, such that *b* = sgn*η* (*c*_2_ – κ|η|)/Δ*t* where sgn *η* denotes the sign of the error angle and |*η*| denotes its magnitude at the moment the steering correction is applied. In cases where this steering correction would bring the pursuer’s flight direction within the error angle tolerance *c*_2_ of another obstacle, the bias command is modified to target the midpoint of the gap between them. This discontinuous open-loop implementation has a clear behavioral interpretation, in that the bird is assumed to avoid obstacles by making a saccadic flight maneuver analogous to those observed in insects.

It is reasonable to suppose that an analogous biased guidance model might successfully describe obstructed pursuit in insects, particularly given the saccadic nature of their flight maneuvers, but previous work on obstructed pursuit in robber flies *Holcocephala fusca* has instead modelled obstacle avoidance as a closed-loop response [13], with smooth turning commanded as:

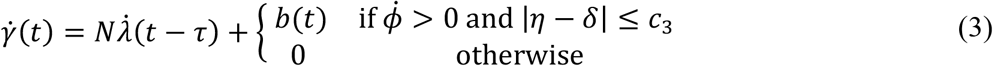

where 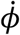 is the looming rate of a narrow object (i.e., the rate of change in its apparent angular width). Here, the guidance constant *N* = 3.6 and delay *τ* = 0.03 s are fitted parameters, whilst *c*_3_ = 43° is the half-width of the region of interest about the target within which looming objects are treated as obstacles. Whilst this is still a discontinuous model of obstacle avoidance, in the sense that the bias command *b* is only engaged under certain conditions, it is implemented by feeding back the looming rate 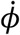 as 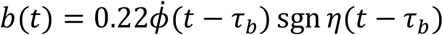 with delay *τ_b_* = 0.09 s. Hence, because 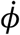 increases exponentially on approach, so too will the bias command *b*, except insofar as it causes the pursuer to turn away from the obstacle. Eq. 3 has some clear disadvantages, in that it would be complex to implement for a dense obstacle field like the one used in our experiments, and commands avoidance of objects that may not necessarily pose a collision risk. It would therefore be of interest to test whether the simpler feedforward model of obstacle avoidance that we have proposed (Eq. 2) can successfully model obstructed pursuit in insects.

## Physiological implementation of biased guidance

How might the steering bias that we have modelled for Harris’ hawks be implemented physiologically? The bias command *b* in Eq. 2 is applied at a distance *d* ≤ 4 m from an upcoming obstacle when κ|*η*| ≤ sin^-1^(0.6/d), although it is probable that the birds would have used optic flow cues to estimate their time to collision with the obstacle rather than its absolute range [18]. Under this model, at the threshold distance of *d* = 4 m (or equivalent time to collision), a steering correction of *b*Δ*t* = 9° – δ|*η*| will be applied if κ|*η*| ≤ 9°. Here *η* is the error angle between the pursuer’s flight direction and its line-of-sight to the near edge of the obstacle, and *κ* = –1 if the pursuer is on a direct collision course with the obstacle, with κ = 1 otherwise. The most direct way of estimating these quantities is from the optic flow field, which is especially straightforward if gaze is stabilized rotationally such that the pursuer’s flight direction coincides with the center of expansion of what is then a pure translational optic flow field. In this case, the condition κ|η| ≤ 9° is met whenever the center of expansion appears either directly on the obstacle (κ = –1), or on the background (κ = 1) within 9° of the edge of the obstacle. Moreover, the error angle *η* is equal to the angle between the center of expansion and the near edge of the obstacle.

In practice, most visually guided pursuers track their target by turning their head, which complicates the interpretation of the optic flow field by combining rotational and translational self-motion components. In an ideal tail-chase, however, the pursuer’s flight direction becomes aligned with the line-of-sight to its target as the deviation angle *δ* is driven towards zero. Hence, another simple heuristic, applicable only in a tail-chase, is to approximate the error angle *η* as the difference in azimuth between the target and the near edge of the obstacle. Moreover, a recent pilot study [17] of Harris’ hawk gaze strategy during obstructed pursuit found that the bird fixated its target at an azimuth of ±10° with respect to the sagittal plane of its head, coinciding with the assumed projection of its left or right temporal fovea. If this anecdotal result generalizes, such that targets are fixated at ±10° on the right (left) temporal fovea when turning to the right (left) around an obstacle, then at the threshold distance of *d* = 4 m, the steering correction *b*Δ*t* = 9° – κ|*η*| that Eq. 2 demands would be approximately the azimuth of the obstacle’s edge with respect to the head’s sagittal plane. Equivalently, if the pursuer’s gaze were shifted to fixate the obstacle’s edge in the head’s sagittal plane, as has been observed in birds [19] including Harris’ hawks [17], then the amplitude of the required body saccade would be approximately the amplitude of the completed head saccade.

## Applications of biased guidance in autonomous systems

The model of obstructed pursuit that we have identified for Harris’ hawks (Eq. 2) is closely related to a form of guidance law used in missile engineering, called biased proportional navigation [20]. This is a modification of the proportional navigation guidance law 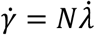 with a bias command *b* added such that 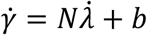, but is often expressed in the alternative form 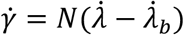, by making the substitution 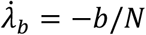. Typically, the bias command *b* is used to modify the agent’s underlying targeting response so as to accomplish some other objective, such as optimizing the control efficiency of a rocket [20], causing a missile to attain a required impact angle [21], guiding an autonomous vehicle along a specified path [22], or meeting specific rendezvous conditions in spaceflight [23]. Many different variants of biased proportional navigation have been proposed, with bias commands that may be engaged in either a continuous or discontinuous fashion, and that may be specified in either open or closed loop [24]. Our modelling demonstrates another possible technical application of biased guidance, where the bias command is used to implement obstacle avoidance in conjunction with target pursuit. This approach differs fundamentally from previous studies that have used unbiased proportional navigation to model collision avoidance in birds [2] or autonomous vehicles [25] by treating the clearance from an object as the target of the proportional navigation guidance law itself (i.e. where 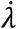 is defined as the line-of-sight rate of the clearance). Biased guidance therefore offers a biologically inspired mechanism for resolving the conflict between obstacle avoidance and target pursuit, which could be deployed in drones designed to intercept other drones in clutter. As the same guidance laws can also be used to steer flight towards stationary targets [2, 16], application of an open-loop bias command could also be used for obstacle avoidance during homing flight, or when flying between waypoints.

## Conclusions

Although it is possible that other guidance laws [15] might explain our hawks’ pursuit behavior as well as the mixed guidance law we have fitted (Eq. 1), our modelling demonstrates high repeatability in the guidance parameters fitted across hundreds of flights collected under varying experimental conditions (Fig. S1B,C), including different studies on different individuals [11]. Such quantitative repeatability is rare in behavioral studies, and presumably reflects both the goal-directed nature of the task and the accuracy of the kinematic measurements used to describe it. In summary, we find that pursuit behavior in Harris’ hawks is well modelled by assuming that their turn rate 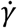 is commanded by feeding back both the angular rate 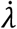 of their line-of-sight to the target, and the deviation angle *δ* between their flight direction and line-of-sight to the target. This target pursuit subsystem serves to drive the pursuer’s deviation angle *δ* to zero, leading to a tail-chase that promotes implicit obstacle avoidance if their target follows a safe path through clutter (Fig. S1A). In addition, we find that Harris’ hawks bias their take-off direction (Fig. 3) and make mid-course steering corrections (Fig. 5) that perturb the deviation angle *δ* when a collision is imminent (Fig. 4), thereby implementing explicit obstacle avoidance (Fig. 6). This obstacle avoidance subsystem is well modelled by assuming that the hawks make a discrete steering correction when they encounter an obstacle blocking their path at close range, aiming for a clearance of just over one wing length from its nearest edge. Harris’ hawks therefore resolve the conflict between obstacle avoidance and prey pursuit by applying an open-loop bias command that modifies their closed-loop targeting response in a discontinuous fashion.

## Methods

### Experimental design

We recorded the flight trajectories of N=4 captive-bred Harris’ hawks *Parabuteo unicinctus* pursuing a falconry lure towed along a zigzagging course around a set of pulleys, with or without obstacles present (Fig. 1A). The birds included one 7-year old female (Ruby) that had been included in a previous study [11], plus three first-year males (Drogon, Rhaegal, Toothless) that had not previously chased a target. A subset of the flights without obstacles that we report are described and analyzed elsewhere using a related method [15], but the flights with obstacles are reported here for the first time. Each bird usually flew after the lure four times per day, taking off spontaneously from the falconer’s gloved fist when the lure began moving. The lure was hidden inside a tunnel at the start of each test, mimicking a terrestrial prey item being flushed from cover. The lure vanished into another tunnel if the bird failed to catch it by the end of the course, which motivated the birds to catch the lure whilst it was still moving.

The experiments began with an 8-day training phase to familiarize the hawks with the task of chasing the lure without obstacles. This yielded a set of n=128 obstacle-free training flights, following which we introduced obstacles into the environment. We conducted a single day of obstacle familiarization flights, using an open layout comprising two rows of four ropes. This yielded a set of n=16 obstacle familiarization flights during which the lure was pulled through the gaps between the obstacles. We used a different obstacle arrangement for the main test flights: the first row of test obstacles comprised an impenetrable grille of eight ropes centered on the midline of the flight hall (Fig. 1B); the second row of test obstacles comprised four pairs of ropes blocking each of the lure’s four possible paths on its way to the last set of pulleys (Fig. 1A). This yielded a total of n=106 obstacle-free test flights and n=154 obstacle test flights, recorded over 15 days of trials including 7 days with obstacles, 5 days without obstacles, and 3 days at the start of the period in which the presence or absence of obstacles was randomized between flights.

We used a simplified pulley configuration at the start of the initial training phase, with four pulleys placed in a diamond-shaped configuration (Pulleys 1-4 in Fig. 1A). This layout produced two possible lure courses, with an unpredictable bifurcation at the first pulley followed by two predictable changes in target direction at the next two pulleys. We modified the pulley setup before the end of the training phase, placing six pulleys in a chevron-shaped configuration (Fig. 1A,B). This layout produced six possible courses, with two or three unpredictable bifurcations in target direction, and one predictable change in direction at the last pulley. The lure course and hawk starting position were randomly assigned before each flight, and we laid dummy towlines to make it harder for the hawks to anticipate the lure’s course (Fig. 1A,B). The speed of the lure was randomized within the range 6-8 m s^-1^ for each flight; at higher speeds, the hawks were unable to catch the lure before the end of the course. Following the initial training phase, we randomized the presence or absence of obstacles between test flights. This took considerable time, however, and was an unnecessary source of stress for the birds, so we subsequently randomized the presence or absence of obstacles once at the start of each day.

### Experimental protocol

The experiments were carried out at the John Krebs Field Station, Wytham, Oxford, UK between January and March 2018 in a windowless flight hall measuring 20.2 m by 6.1 m, with an eaves-height of 3.8 m. The flight hall was lit by flicker-free LED up-lights providing approximately 1000 lux of diffuse overhead lighting reflected by white fabric sheets hung from the ceiling to mimic overcast morning or evening conditions. The walls of the hall were hung with camouflage netting to provide visual contrast, and small shrubs and trees were placed down the sides of the room to discourage flight outside of the central test area (Fig. 1B). The hawks were flown individually from the gloved fist of a falconer positioned in one of three starting positions across the flight hall (Fig. 1A). A falconry lure with a small food reward attached was towed around a series of large pulleys by two Aerotech linear actuators rigged with a block and tackle system to increase their output speed (ACT140DL, Aerotech Limited, Hampshire, UK); a drag line pulled along behind the lure smoothed its path around the pulleys (Fig. 1A). For the experiments with obstacles, we hung jute ropes (diameter: 0.05 m) from the roof space to the floor to mimic compliant stems or branches, wrapping them in expanded polystyrene pipe insulation to make them safe in case of collision (Fig. 1B).

We reconstructed each flight using 20 motion capture cameras recording at 200 Hz (Vantage 16, Vicon Motion Systems Ltd, Oxford, UK), under stroboscopic 850 nm infrared illumination outside the visible spectrum of Harris’ hawks [26]. Four high-definition video cameras (Vue, Vicon Motion Systems Ltd, Oxford, UK) recorded synchronized reference video at 120 Hz. The cameras were mounted on a scaffold at a height of 3 m, spaced around the perimeter of the flight hall to maximize coverage (Fig. 1A,B). The motion capture system was turned on at least an hour before commencement of the flight experiments and was calibrated immediately before the first trial by moving an Active Calibration Wand (Vicon Motion Systems Ltd, Oxford, UK) through the capture volume. The origin and ground plane of the coordinate system were set by placing the calibration wand on the floor in the center of the room. Each bird was fitted with two rigid marker templates (Fig. 1C): a backpack template with four 6.4 mm diameter spherical retroreflective markers arranged in an asymmetric pattern, attached to a falconry harness (Trackpack Mounting System, Marshall Radio Telemetry Ltd, Cumbria, UK); and a tail-pack with three 6.4 mm diameter retroreflective markers, attached to a falconry tail mount (Marshall Aluminium Tail Feather Piece, Marshall Radio Telemetry Ltd, Cumbria, UK). The birds also wore retroreflective markers attached directly to the feathers on their head, wings, or tail, but these are not included in the present analysis. Six 6.4 mm diameter retroreflective markers were attached directly to the lure, with three markers on either side in a back-to-back arrangement. Each rope obstacle was fitted with 9.5 mm diameter markers at eye level and floor level.

### Trajectory reconstruction

The three-dimensional (3D) positions of the bird, lure, and obstacle markers were reconstructed using Nexus software (Vicon Motion Systems Ltd, Oxford, UK), in a coordinate system aligned to the principal axes of the flight laboratory. Previous work had found that the Vicon software was not always able to identify which marker was which between frames, owing to marker occlusion and the small distance between the markers relative to the distance travelled between frames [27]. We therefore used custom-written code in Matlab (Mathworks Inc, MA, USA) to label the anonymous markers in the rigid templates. Our first step was to identify markers that remained stationary through the trial as being obstacle markers. For the remaining markers, we used their height above the floor to distinguish between markers on the bird and the lure and used a clustering algorithm to distinguish between markers on the backpack and the tail-pack. We used the centroid of the backpack and lure as our initial estimate of their respective positions, treating any frames in which fewer than three markers were detected on the backpack, tail-pack, or lure as missing data.

The initial position estimates for the backpack, tail-pack and lure were contaminated by misidentified markers, which we excluded by removing points falling further than 0.5 m from the smoothed trajectory obtained using a sliding window mean of 0.05 s span. We then repeated this sliding window mean elimination on the raw data with extreme outliers excluded, this time using a distance threshold of 0.075 m. Our next step was to crop the trajectories to begin at the first frame on which both the bird and lure were visible, and to end at the point of intercept defined as the point of minimum distance between the bird and lure. We then used cubic interpolation to fill in any missing data points and fitted a quintic spline to smooth the 3D data, using a tolerance of 0.03 m for the bird and 0.01 m for the lure. Finally, we double-differentiated the spline functions, which we evaluated analytically to estimate the velocity and acceleration of the bird and lure at 20 kHz, resulting in a suitably small integration step size for our simulations.

### Guidance simulations

As the birds always flew close to the ground plane, our guidance analysis concerns only the horizontal components of the pursuit. We used the same forward Euler method and Matlab code described previously [11] to simulate the hawk’s horizontal flight trajectory given the measured trajectory of the lure. We modelled the hawk’s turning using the mixed guidance law in Eq. 1 for a given set of parameter settings *N, K*, and *τ*, matching its simulated flight speed to its measured flight speed. In cases where the hawk’s simulated trajectory resulted in an earlier intercept than its measured trajectory, we matched the continuation of the simulated trajectory to that of the lure up to the measured point of intercept. By default, we matched the hawk’s initial flight direction in the simulations to that which we had measured. However, we also ran versions of the simulations in which we re-initialized the hawk’s flight direction at take-off or 4 m from the second obstacle, by directing its flight towards some specified location (see Results). We defined the prediction error for each flight, *ε*(*t*), as the distance between the measured and simulated flight trajectories.

### Statistical analysis

We optimized the guidance parameters *N*, *K*, and *τ* by minimizing the median of the mean prediction error, 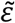, over a given subset of flights. We did this using an exhaustive search procedure for values of *N* and *K* from 0 to 2 at intervals of 0.05, and for values of *τ* from 0 to 0.09 s in intervals of 0.005 s. To ensure that we modelled the same section of flight for all values of τ, we began each simulation at 0.09 s after the start of the trajectory. Although we optimized the guidance parameters for the obstacle and obstacle-free test flights separately at first, we subsequently combined these subsets, owing to the observed similarity of their best-fitting parameter settings. Because there were more test flights with obstacles than without, we used a balanced subsampling procedure to avoid biasing the fitting of the joint model in favor of obstructed pursuit. Specifically, we sampled 80 flights at random from each subset and identified the parameter settings that minimized 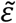 over that sample. We repeated this sampling experiment 100,000 times and took the grand median of the resulting best-fitting parameter settings as our refined model. We quantified the goodness of fit of a given guidance model by computing the mean prediction error, 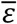, for each flight. We then used a bias corrected and accelerated percentile method to compute a bootstrapped 95% confidence interval for the median of the mean prediction error 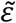 at the best-fitting parameter settings. We report bootstrapped 95% confidence intervals for other properties of the flight trajectories where relevant.

### Ethics statement

This work was approved by the Animal Welfare and Ethical Review Board of the Department of Zoology, University of Oxford, in accordance with University policy on the use of protected animals for scientific research, permit no. APA/1/5/ZOO/NASPA, and was considered not to pose any significant risk of causing pain, suffering, damage or lasting harm to the animals.

### Data availability

Supporting data are available at https://doi.org/10.6084/m9.figshare.21905211.

## Acknowledgments

We thank our falconers Mark Parker, Helen Sanders, and Lucy Larkman for their involvement in the experiments, and our mechanical workshop technician John Hogg for designing and building the equipment. We thank James Shelton and Natalia Pérez-Campanero Antolín for helpful conversations.

## Funding

This project has received funding from the European Research Council (ERC) under the European Union’s Horizon 2020 research and innovation programme (Grant Agreement No. 682501). JK was supported by a Christopher Welch Scholarship from the University of Oxford. LF and SM were supported by funding from the Biotechnology and Biological Sciences Research Council (BBSRC) [grant number BB/M011224/1], via the Interdisciplinary Bioscience Doctoral Training Partnership. For the purpose of Open Access, the author has applied a CC-BY public copyright licence to any Author Accepted Manuscript (AAM) version arising from this submission.

## Author contributions

Conceptualization: CHB, JAK, GKT. Formal analysis: CHB, JAK, LAF, MKH, SM, GKT. Methodology: CHB, JAK, GKT. Investigation: CHB, JAK, MKH, SM. Visualization: CHB. Supervision: GKT. Writing—original draft: CHB. Writing—review & editing: GKT. All authors reviewed and commented on the original draft.

## Competing interests

The authors declare that they have no competing interests.

## Extended Data

**Figure S1.**
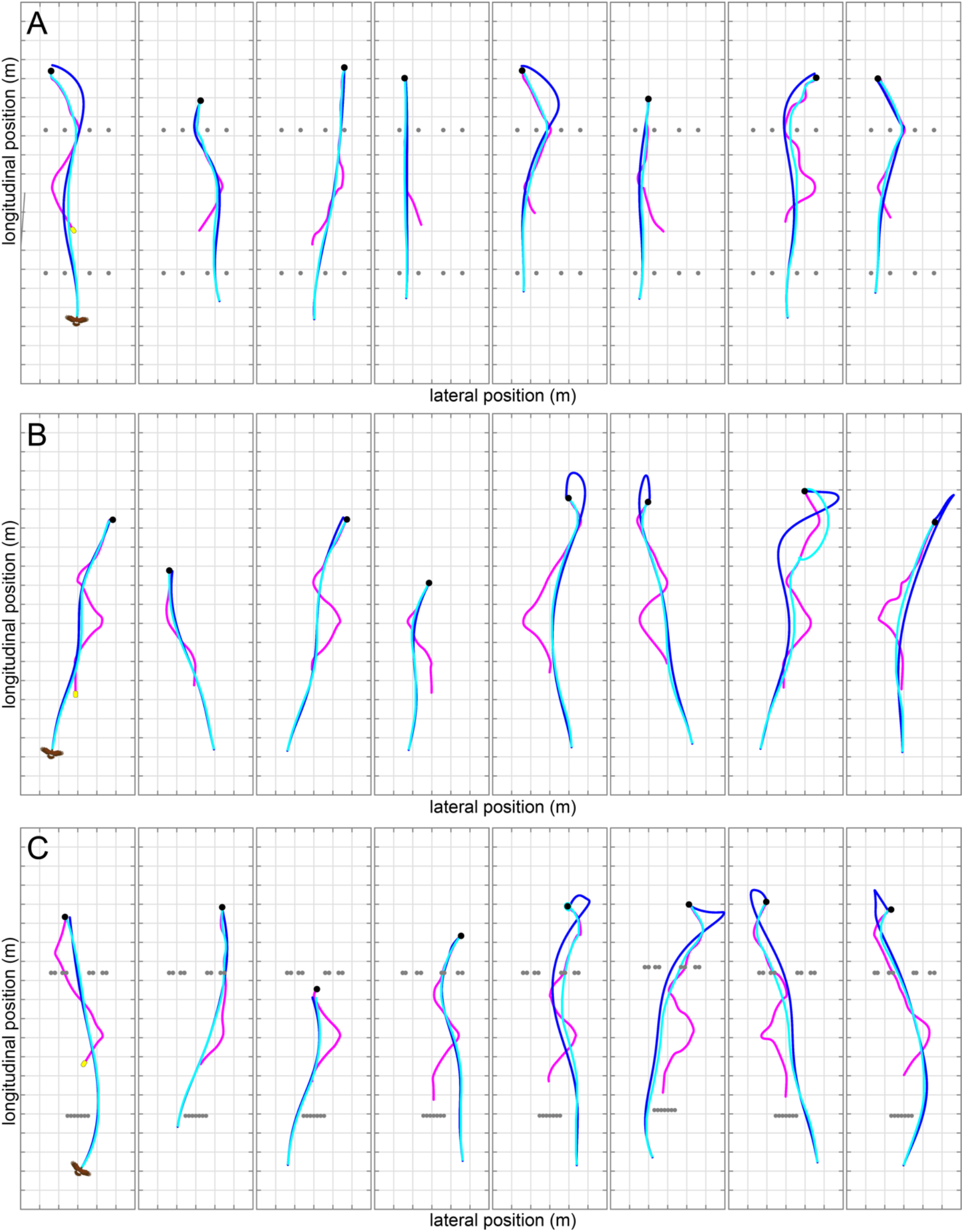
Measured pursuit trajectories of Harris’ hawks compared to guidance simulations under the mixed guidance law. Each panel represents a single flight and plots the hawk’s measured flight trajectory (blue line) in pursuit of the lure (magenta line) up to the point of capture (black dot). The measured data are compared to a simulation of the hawk’s trajectory (cyan line) under the mixed guidance law (Eq. 1). Hanging rope obstacles are plotted as grey dots if present. Grid spacing: 1 m. **(A)** Guidance simulations inheriting the parameter settings *N* = 0.7, *K* = 1.2 s^-1^ and *τ* = 0.09 s fitted previously [28], shown for the eight longest obstacle familiarization flights. Note that the lure passes between the obstacles of the second row during these flights, and that the tail-chasing behavior which the mixed guidance law promotes leads to implicit obstacle avoidance as a result. **(B,C)** Guidance simulations under the refined mixed guidance law with best-fitting parameters *N* = 0.75, *K* = 1.25 s^-1^ and *τ* = 0.01 s fitted jointly to the n=106 obstacle-free test flights (B) and the n=154 obstacle test flights (C). Panels on the left show the four flights with the lowest mean prediction error relative to the total distance flown, for flights > 9 m in length. Panels on the right show the four longest flights. Grid spacing: 1 m.

## Notes

https://doi.org/10.6084/m9.figshare.21905211

### Competing Interest Statement

The authors have declared no competing interest.

### Summary of Updates

Updated main text and summary, and moved one figure to Supplementary Information.

